# CiteFuse enables multi-modal analysis of CITE-seq data

**DOI:** 10.1101/854299

**Authors:** Hani Jieun Kim, Yingxin Lin, Thomas A. Geddes, Jean Yang, Pengyi Yang

## Abstract

Multi-modal profiling of single cells represents one of the latest technological advancements in molecular biology. Among various single-cell multi-modal strategies, cellular indexing of transcriptomes and epitopes by sequencing (CITE-seq) allows simultaneous quantification of two distinct species: RNA and surface marker proteins (ADT). Here, we introduce CiteFuse, a streamlined package consisting of a suite of tools for pre-processing, modality integration, clustering, differential RNA and ADT expression analysis, ADT evaluation, ligand-receptor interaction analysis, and interactive web-based visualization of CITE-seq data. We show the capacity of CiteFuse to integrate the two data modalities and its relative advantage against data generated from single modality profiling. Furthermore, we illustrate the pre-processing steps in CiteFuse and in particular a novel doublet detection method based on a combined index of cell hashing and transcriptome data. Collectively, we demonstrate the utility and effectiveness of CiteFuse for the integrative analysis of transcriptome and epitope profiles from CITE-seq data.

## Introduction

The latest advancement in multi-modal profiling of single cells promises to revolutionise our understanding in cellular biology that was previously inconceivable through bulk profiling technologies (Datlinger et al., 2017; Macaulay et al., 2015; Mohammed et al., 2017). Among various single-cell multi-modal strategies, cellular indexing of transcriptomes and epitopes by sequencing (CITE-seq) (Stoeckius et al., 2017) and its variants such as RNA expression and protein sequencing (REAP-seq) (Peterson et al., 2017) represent a class of approaches that allows simultaneous quantification of global gene expression and cellular proteins using single-cell RNA-sequencing (scRNA-seq) and antibody-derived tags (ADTs), respectively, on single cells. Further extensions such as multiplexed detection of proteins, transcriptomes, clonotypes and CRISPR perturbations enable additional modalities to be profiled on single cells (Mimitou et al., 2019).

While the surface proteins of individual cells measured by ADTs are also transcriptomically profiled by scRNA-seq, the measurements of these two different molecule species produced from the same genes do not necessarily correlate with each other, presumably because of post-transcriptional and post-translational gene regulation (See, Lum, Chen, & Ginhoux, 2018). Therefore, computational integration of single cell multi-modal profiling data may allow a more accurate characterisation of cells (e.g., cell type identification) (Buettner et al., 2015) and provide new biological insights that may be observable from neither a single data source (Lin et al., 2019) nor modality (Stuart et al., 2019).

Here we present CiteFuse, a computational framework that implements a suite of methods and tools for CITE-seq data from pre-processing through to integrative analytics. This includes doublet detection, network-based modality integration, cell type clustering, differential RNA and ADT expression analysis, ADT evaluation, ligand-receptor interaction analysis, and interactive web-based visualisation of the analyses (**Figure 1A**). Using both simulations and an experimental CITE-seq dataset generated from PBMCs (Mimitou et al., 2019), we demonstrate the integrative capacity of CiteFuse in various scenarios and its advantage over analysing each individual source and modality of data. CiteFuse represents the first method specifically designed to systematically integrate RNA and ADT modalities of single cells in CITE-seq data. We anticipate its increasing utility given the rapidly accumulating volume of multi-omic and multi-modality single cell data generated using CITE-seq from various biological studies (Mimitou et al., 2019; Stoeckius et al., 2017). Finally, CiteFuse is implemented as an R package (http://SydneyBioX.github.io/CiteFuse/) as well as a user-friendly web application (http://shiny.maths.usyd.edu.au/CiteFuse/), allowing users to upload and analyse their CITE-seq datasets.

**Figure 1.**
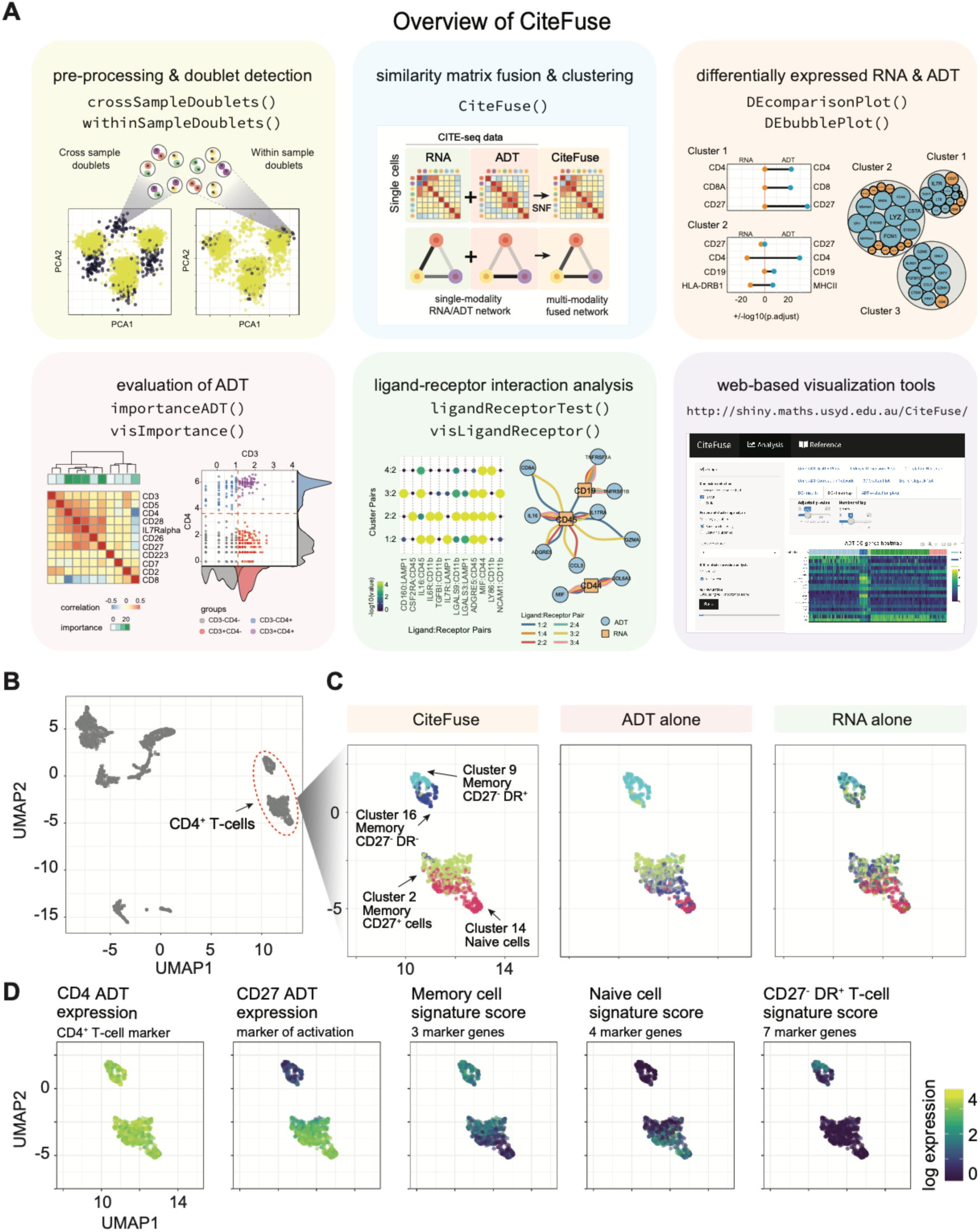
An overview of CiteFuse and application to clustering of PBMC CITE-seq data. (A) A summary of the key components and functions implemented in CiteFuse. (B) UMAP visualisation of human PBMC CITE-seq data (Mimitou et al., 2019). (C) Clustering outputs (represented by colours of points) of CD4+ T-cells using multi-modality (CiteFuse), or single-modality (antibody-derived tag [ADT] or RNA alone). (D) Expression of key markers of sub-cell types in CD4+ T-cells.

## Results

### CiteFuse gains information from multi-modal integration of CITE-seq data

To take advantage of the complementary information present in multi-modal CITE-seq data, CiteFuse integrates mRNA and ADT expression by constructing networks across single cells for each data modality and fusing these networks using a similarity network fusion algorithm (Wang et al., 2014) (**Figure 1A, blue tile**). It subsequently uses a spectral clustering algorithm to cluster the cells based on the fused matrix. To test whether there is any advantage in using the fused multi-modal expression matrix over the single-modal matrices, we performed a comparison between the different modalities and across existing clustering algorithms with simulated CITE-seq data (Zhang et al., 2019) (**Figure S1A**). We demonstrate that in both “easy” and “hard” scenarios (see Methods), CiteFuse clusters cells more accurately than directly applying spectral clustering on the two single-modal data types (**Figure S1B**). Moreover, we demonstrate that CiteFuse performs better compared to several established clustering procedures, including SIMLR (Wang et al., 2017), PCA + *k*-means, and Seurat (Satija et al., 2015) with either RNA or ADT expression matrix (**Figure S1B**).

To test if the information gain from multi-modal analysis using CiteFuse observed from the simulation study translates into real-world data analysis, we next applied CiteFuse to a recent human PBMC CITE-seq dataset (Mimitou et al., 2019) (**Figure 1B**). We show that clustering using CiteFuse on multi-modal data and directly applying spectral clustering on single-modal (ADT or RNA) data lead to different clustering outcomes (**Figure S2A**). We found that CiteFuse can generate four CD4+ T-cell clusters (**Figures 1C and S2B**), of which three are CD4+ memory T-cells (clusters 2, 9, and 16) expressing high level of S100A4 (a marker of memory T-cells) and one is CD4+ naive T-cells (cluster 14) expressing high level of SELL (a marker of naive T-cells) (Elyahu et al., 2019; Haining et al., 2008) (**Figure S2C**). In contrast, clustering using ADT alone leads to over-partitioning of CD4+ T-cells into five clusters and clustering using RNA alone leads to under-partitioning of these cells into three clusters (**Figures S2B and S2C**).

Moreover, we observed that clustering using RNA-alone fails to partition CD27+ and CD27-populations of memory T-cells, whilst clustering using CiteFuse or ADT-alone can discriminate these two populations, albeit to different resolutions (**Figures 1D and S2B**). A closer examination of the CD27-CD4+ memory T-cell subpopulations (**Figure 1C**; light and dark blue clusters in CiteFuse; light blue cluster in ADT alone) reveals that only CiteFuse can discriminate between CD27-DR+ (light blue) and CD27-DR-(dark blue) memory T-cell subpopulations (Fonseka et al., 2018) (**Figures 1D and S2D**), revealing that only CiteFuse has the capacity to finely map T-cell subpopulations and further demonstrates the gain in information CiteFuse benefits from multi-modal analysis.

### CiteFuse detects both cross- and within-sample doublets

Identification and removal of doublets from scRNA-seq data derived from microfluidic technology is essential for downstream analysis. Cell hashing is a multiplexing technique commonly used in CITE-seq for pooling multiple samples (Stoeckius et al., 2018). Because a key principle in cell hashing is the selection of ubiquitously and highly expressed surface markers, against which distinct hashtag oligonucleotide (HTO)-conjugated antibodies are raised, the high number of the ubiquitous epitopes raises the possibility of utilising HTO-derived expression to detect within-sample doublets marked by anomalous HTO expression. To this end, CiteFuse takes advantage of the matched matrices for RNA, ADT, and HTO expression generated from a CITE-seq experiment (**Figure 2A**) and implements a stepwise approach to detect and filter both cross- and within-sample doublets (**Figure 2B**). In the first step, a Gaussian mixture model is used to identify cross-sample doublets that have more than one hashtag (i.e. stained by orthogonal HTOs) (**Figure S3A**). Next, by leveraging the ubiquitous nature of HTO expression, CiteFuse detects within-sample doublets from DBSCAN clustering of single cells based on two features—total number of captured unique molecular identifiers (UMIs) and total HTO expression (**Figure 2B**). Data are filtered in step one based on the mixture modelling step for cross-sample doublets and then based on a baseline HTO threshold calculated through the Gaussian mixture model for within-sample doublets (see Methods).

**Figure 2.**
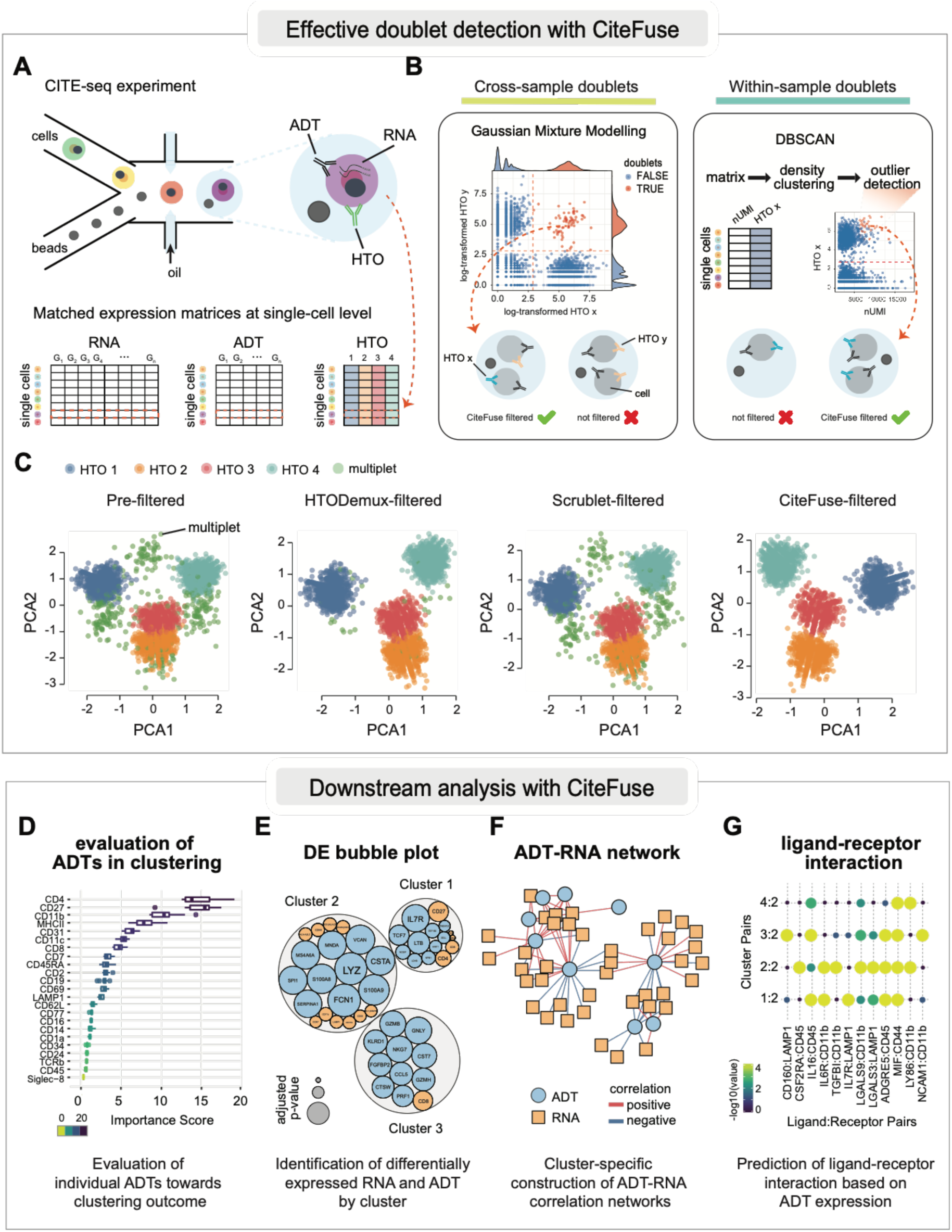
Doublet detection and downstream analysis using CiteFuse. (A) A schematic representation of the CITE-seq experiment and the cell hashing data generated by using hashtag oligonucleotide (HTO). (B) The doublet detection approach implemented in CiteFuse. This includes cross-sample doublet identification using Gaussian mixture modelling and a novel within-sample doublet identification method using a combined index of cell hashing and transcriptome data. (C) PCA visualisation of HTO expression before and after filtering of doublets using HTODemux (Stoeckius et al., 2018), Scrublet (Wolock et al., 2019), or CiteFuse. (D-G) Key downstream analytical tools implemented in CiteFuse.

We benchmarked our doublet filtering approach with alternative methods, HTODemux (Stoeckius et al., 2018) and Scrublet (Wolock et al., 2019), on the PBMC dataset (Mimitou et al., 2019) by demonstrating that doublets/multiplets detected through CiteFuse show comparably high number of unique genes and UMIs (**Figure S3B**). Notably, we show that the within-sample doublets identified through CiteFuse represent outlier cells that have both high total UMIs and high HTO expression (**Figure S3C**). We show that our approach captures most doublets detected through HTODemux and Scrublet but also identifies additional ones that may have been missed by HTODemux and Scrublet (**Figure S3D**). When we quantified the total UMIs and number of unique genes in cells exclusively identified by each method (**Figure S3E**), we found that doublets exclusively detected by HTODemux and Scrublet show characteristics that resemble singlets whereas those only detected by CiteFuse resemble doublets (**Figures S3B and S3E**).

Strikingly, we observed the most improved separation of clusters on the first two principal components of HTO expression before and after filtering of doublets detected by CiteFuse (**Figure 2C)**, suggesting our CiteFuse pipeline enables more accurate filtering of both within- and cross-sample doublets when HTO libraries are available.

### CiteFuse doublet filtering preserves the separation between T-cell subpopulations

To evaluate the impact of filtering method on the downstream analysis, we applied CiteFuse clustering on data either unfiltered (4292 cells) or filtered using the different doublet detection methods—HTODemux (3753 cells), Scrublet (3968 cells), and CiteFuse (3612 cells). Visualisation of the clusters on UMAP revealed very different clustering outcomes by each filtering method, revealing that filtering method can have a large impact on downstream analysis (**Figure S4A**).

We demonstrate the impact of filtering method on downstream analysis by evaluating the capacity of the unfiltered and filtered datasets to define CD4+ and CD8+ T-cell types, two major groups of T lymphocytes. We found that the CiteFuse-filtered dataset leads to the best separation of CD4+ (clusters 2, 9, 14, and 16) and CD8+ (clusters 3, 7, 10, and 15) T-cell populations on the basis of purity scores (**Figure S4B**). Moreover, our results showed that the CiteFuse-filtered dataset can further discriminate CD27+ and CD27-subpopulations within CD4+ and CD8+ T-cells (**Figure S4C-E**). Surprisingly, we observed that HTODemux- and Scrublet-filtered datasets have low capacity to discriminate between CD4+ and CD8+ T-cells, let alone CD27+ and CD27-subpopulations within each of the major T-cell populations (**Figure S4C-E**).

### CiteFuse enables the evaluation of ADTs and visualisation of ADT-RNA networks

The selection of a set of ADTs for CITE-seq may be an expensive process, requiring in many cases optimisation through flow cytometry for antibody concentration and selection. To maximise the selection of ADTs for subsequent CITE-seq experiments, CiteFuse implements a set of evaluation tools that enables CITE-seq end-users to assess ADTs for relative importance and potential redundancy (**Figure 2D**). This includes correlating and visualising ADTs based on their expressions (**Figure S5A**) as well as calculating the relative importance of individual ADTs based on CiteFuse clustering outcome using a random forest model (see Methods) (**Figure S5B**). For example, in the PBMC CITE-seq dataset, we found that CD223 and IgG1 are the two ADTs receiving the lowest importance scores and therefore may not provide much additional information for cell type clustering. Indeed, we observed minimum changes in the clustering outcome (ARI=0.99) even without the two ADTs (**Figure S5C**). We find that more ADTs can be excluded (Subsets 2-3) with minimal effect on clustering results. In addition to ADT evaluation, CiteFuse can also perform cluster-specific differential gene expression analysis to detect and compare differentially expressed RNA and ADT (**Figure 2E**) and generate visualisation of ADT-RNA correlation networks unique to each cluster, allowing users to evaluate relationships between ADT and RNA in an intra-cluster manner (**Figure 2F**).

### CiteFuse facilitates accurate identification of ligand-receptor interactions

Most studies on ligand-receptor interaction in single-cell biology rely solely on mRNA expression (Vento-Tormo et al., 2018), thereby making an implicit assumption that the level of mRNA expression is a proxy for the cell-surface protein expression. Yet studies have shown that the levels of mRNA and proteins of the same gene can vary widely (Gry et al., 2009; Liu, Beyer, & Aebersold, 2016). In case of cell-surface proteins, this is further complicated by the amount of proteins translocated to plasma membrane. CITE-seq opens the possibility to use protein expression at the cell-surface to predict ligand-receptor interactions. To this end, we predicted ligand-receptor interactions based on mRNA expression of the ligand and ADT expression of the receptor, after normalisation and scaling of the mRNA and ADT expression data (see Methods) (**Figures 2G and S6A**). We compared the ligand-receptor interactions identified by CiteFuse with those identified from the conventional approach where the expression of RNA alone is used as a readout for both ligand and receptor expression (**Figure S6A**). We found that the overlap in interactions between the conventional approach and CiteFuse was variable across clusters, but generally a large portion of the ligand-receptor interactions identified through the conventional approach (referred to as RNA-specific) were not identified as interactions through CiteFuse (**Figure S6B**). We also observed in each cluster a fraction of interactions that were identified only by CiteFuse (referred to as CiteFuse-specific) (**Figure S6B**).

We then hypothesised that the large proportion of interactions in the conventional approach that are not detected by CiteFuse may be because of false positive predictions. To investigate this, we calculated the normalised log expression of the ADT and mRNA of all receptors that were identified in a ligand-receptor interaction for each category (CiteFuse-specific, RNA-specific, and Common). We found that although the mRNA expression of the receptors was comparable between the categories the ADT expression of these receptors was much lower in the RNA-specific group than the other two groups (**Figure S6C**). Notably, we found that a strong positive correlation of ADT and mRNA expression (ranked relative to each cluster; see Methods) for receptors identified in a ligand-receptor interaction in the Common and CiteFuse-specific categories but no correlation for those in the RNA-specific category (**Figure S6D**). Similarly, we show that the mRNA expression of ligands detected in the RNA-specific category have higher rankings than those detected in the other two categories (**Figure S6E**). These data show that interactions identified through the conventional approach, which relies on RNA expression alone, may introduce false interactions. These false interactions may potentially be driven by high RNA expression that is not reciprocated in the cell-surface protein expression and thus demonstrates the need to utilise both mRNA and ADT expression in ligand-receptor interaction predictions (**Figure S6F**).

## Methods

### Integration of CITE-seq data through similarity network fusion and spectral clustering

To integrate multi-modal CITE-seq data, CiteFuse first normalises the ADT expression through centred log-ratio (CLR) transformation. It next calculates cell-to-cell similarity matrices from ADT expression using *perb* similarity metric from the *propr* package (Quinn, Richardson, Lovell, & Crowley, 2017) and RNA expression using Pearson’s correlation on highly variable genes identified with the *scran* package (Lun, McCarthy, & Marioni, 2016). The two similarity matrices are scaled using an exponential similarity kernel and then fused by a similarity network fusion algorithm (Wang et al., 2014).

CiteFuse performs spectral clustering (Ng, Jordan, & Weiss, 2002) to identify clusters from the fused similarity matrix. Spectral clustering on single-modal matrices from CITE-seq data were performed for comparison. As well as spectral clustering, CiteFuse also provides the additional option of Louvain clustering (Blondel, Guillaume, Lambiotte, & Lefebvre, 2008), which is an algorithm based on the shared nearest neighbours, which CiteFuse identifies from the fused similarity matrix. Finally, UMAP or tSNE can be applied to the fused similarity matrix to visualise the multi-modal data.

### CITE-seq data simulation and evaluation of CiteFuse

To evaluate the integrative capacity of CiteFuse, we simulated CITE-seq data with SymSim (Zhang, Xu, & Yosef, 2019) and assessed the difference in clustering outcome between the modality of data and also by different clustering methods.

For each simulation, we generated a dataset of 500 single cells among which were six cell types where total numbers of RNA and ADT were 10,000 and 100, respectively. The following parameter settings for sigma (σ), which controls within-population variability, and minimum population size (min_pop) were used to simulate CITE-seq data of different levels of difficulty.

- Simulation 1 (easy): σ (RNA) = 0.8; σ (ADT) = 0.2; and min_pop = 50
- Simulation 2 (hard): σ (RNA) = 0.9; σ (ADT) = 0.4; and min_pop = 20

We generated 10 datasets for each simulation setting and benchmarked CiteFuse against spectral clustering on single-modal matrices and three different clustering methods: *k*-means clustering on PCA reduced dimension (PCA + *k*-means), SIMLR (Wang, Zhu, Pierson, Ramazzotti, & Batzoglou, 2017) and Seurat (Satija, Farrell, Gennert, Schier, & Regev, 2015). For PCA + *k*-means, *k*-means clustering was performed on the first 10 principal components. For *k*-means clustering and SIMLR, the number of clusters was set as six so to be consistent with the simulation set-up. While for Seurat, we set the resolution parameters between 1.5 and 2 such that the number of communities detected by Louvain clustering is consistent with the number of cell types in the simulations. The concordance in clustering outcome was evaluated as the adjusted rand index (ARI), where a higher index indicates better clustering performance.

### CITE-seq data from healthy human PBMCs

To demonstrate our method, we used the recently published CITE-seq data (Mimitou et al., 2019). Specifically, we used the ECCITE-seq dataset from PBMC samples isolated from the blood of healthy human controls. The samples from the human healthy PBMC datasets were pooled from 10x libraries with four distinct barcodes, representing the four hashtag oligonucleotides (HTO) used in the cell hashing.

### Calculation of signature scores for T-cell subpopulations

To calculate the signature scores for the various immune populations, we averaged the expression of the following sets of genes that were previously defined as marker genes for the respective cell types of interest:

1. S100A4, CRIP1, and AHNAK were used to define memory CD4+ T-cells (Elyahu et al., 2019; Haining et al., 2008);
2. TCF, ID3, CCR7, and SELL were used to define naive CD4+ T-cells (Elyahu et al., 2019; Haining et al., 2008);
3. GNLY, GZMB, PRF1, GZMA, NKG7, HLA-DRB1, and HLA-DPA1 were used to define CD4+ CD27-DR+ T-cells (Fonseka et al., 2018);

### CiteFuse doublet detection approach

CiteFuse implements a stepwise procedure to identify both the cross-sample doublets and within-sample doublets from CITE-seq data when cell hashing data is available.

1. Cross-sample doublet identification First, we fit a two-component Gaussian mixture model to each log-transformed HTO expression. The intersection point defined from the mixture model is used to categorise each cell in terms of whether the HTO is either highly or lowly expressed. The cells found to have a single highly expressed HTO are considered as singlets whilst those that have two or more highly expressed HTOs are considered as doublets or multiplets. Cells without any highly expressed HTOs are considered as empty droplets.
2. Within-sample doublet identification Data filtered by cross-sample doublets are next subject to within-sample doublet identification using a density-based spatial clustering and noise detection algorithm (DBSCAN) on an HTO-specific matrix comprising of two features—total number of UMIs and log-transformed HTO expression. The two parameters used in the DBSCAN for this study are eps = 190 and minPts = 50. This procedure is repeated for each HTO and the smallest cluster from DBSCAN clustering is assigned as within-sample doublets.

We benchmarked our doublet detection method against two existing methods: HTODemux (Stoeckius et al., 2018) from the Seurat package and Scrublet (Wolock, Lopez, & Klein, 2019). We used the default parameter settings for sim_doublet_ratio and n_neighbors to construct the KNN classifier to simulate doublets with Scrublet by following their online tutorial (https://github.com/AllonKleinLab/scrublet/) and set an expected doublet rate of 0.04. We compared the total number of UMI and the number of unique expressed genes for each cell by each method (HTODemux, Scrublet, and CiteFuse). To compare the effect of filtering method on the downstream analysis, we performed spectral clustering on the output of the similarity network fusion and calculated the purity score of CD8+ cells against CD4+ cells in individual clusters for each filtering method.

### Calculation of purity score

To calculate the purity of CD4+ and CD8+ T-cell populations, we first identified CD4+ and CD8+ T-cells by creating a Gaussian mixture model on expression of CD4, CD8, and CD11c. For CD4+ T-cells, we created a Gaussian mixture model of CD4 and CD11c expression to define the CD4+ CD11c-population. For CD8+ T-cells, the same approach was employed but with CD8 and CD11c expression. Using the threshold calculated from the mixture model, cells were assigned as either CD4+ negative or positive cells and CD8+ negative or positive cells. Next, using the CD4+ and CD8+ T-cell labels, we calculated the purity of each cluster for either CD4 or CD8 T-cells. A purity of 1 denotes a cluster composed purely of either CD4 or CD8 T-cells, and a purity score of 0 denotes a cluster devoid of either cell type.

### Analysis and visualisation of differentially expressed RNA and ADT

To identify the differentially expressed mRNA and ADTs for each cluster, we used the Wilcoxon rank sum test to compare the log-transformed expression of mRNA and ADT for each cluster against all other clusters. The p-values were adjusted using the Benjamini and Hochberg method (Benjamini & Hochberg, 1995).

For the selection of RNA and ADT markers for a given cluster, we considered the following three criteria:

1. An adjusted *P*-value of lower than 0.05;
2. The mean expression of RNA and ADT in the cells of the cluster is greater than the mean expression of RNA and ADT in cells of all other clusters; and
3. The proportion of cells in the cluster expressing the RNA and ADT is greater than the proportion of cells expressing the RNA and ADT across all other clusters by at least 10%.

CiteFuse enables two exploration methods to visualise the results of differential expression analysis for both RNA and ADT in a single plot:

1. DEcomparisonPlot The DEcomparisonPlot visualises the positive log10 transformed adjusted P-values as a dot of the RNA and the negative log10 transformed adjusted p-values of its corresponding ADT signal on the same y-axis.
2. DEbubblePlot We used the circlepack plot to visualise the RNA and ADT markers, where each marker is represented by a circle and the size of the circle represents the magnitude of the negative log10 *P*-value. The circles representative of RNA and ADT markers from the same clusters are then grouped into a larger circle, representing individual clusters. The circlepack plots are generated using the R package *ggraph* (Pedersen, 2017).

### ADT-RNA correlation network construction

To construct the ADT-RNA co-expression network, we calculated the Pearson’s correlation between mRNA and ADT expression. Other correlation calculation methods, such as the Spearman and Kendall correlation, are also available as options in our CiteFuse package. ADT-RNA pairs with high absolute correlation (above a default setting of 0.6) are used to construct the ADT-RNA correlation network. The networks are visualised using R packages, igraph (Csardi & Nepusz, 2006) and visNetwork (Almende & Thieurmel, 2016).

### Evaluation of ADT importance

To evaluate the importance for each ADT towards the clustering outcome, we trained a random forest model on a subset of randomly sampled cells (80% of total), using the clustering labels from the similarity network fusion of the PBMC CITE-seq data. After 50 repeated fitting of the random forest model, we quantified the feature importance in terms of the mean decrease in Gini index as a surrogate of the importance of each ADT towards clustering outcome. We defined ADT importance score as the median of the feature importance of all runs. A higher score indicates greater importance of the ADT.

Next, to identify potentially redundant ADTs that do not contribute significantly towards clustering outcome, we sorted the ADTs by importance and drew cut-offs in accordance to the local maximums of the difference in importance scores. We then retained the subset of ADTs the with importance scores greater than the cut-offs and performed similarity network fusion analysis. We calculated the adjusted rand index (ARI) to measure the concordance in clustering outcome for each subset of ADTs against that of the full dataset.

### Ligand-receptor interaction prediction

One of the key challenges in analysing ligand-receptor relationships between two modalities is the difference in scaling and distribution. To address this, we first scaled each feature into a range of 0 to 1 through min-max normalisation. Specifically, for every value of a feature *x* across all single cells, the normalised expression z is calculated by

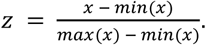

Another challenge we encountered was the difference in distribution between the two modalities: we observed that the distribution of mRNA expression tends to be more zero-inflated than ADT expression. Because comparing unequal distributions has the potential to introduce bias, especially during ligand-receptor predictions when the mean expression is compared, we thus performed another step of transformation on the ADT expression to force the low-expression values to zero. For the normalised expression *z*, with *z* ∈ [0,1], the transformed expression is calculated by

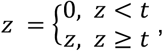

where *t* is set as 0.5 by default.

Lastly, we performed a similar procedure to the method from Vento-Tormo et al. to predict ligand-receptor interactions (Efremova, Vento-Tormo, Teichmann, & Vento-Tormo, 2019). For each ligand-receptor interaction pair originating from a cluster expressing the ligand and another cluster expressing the receptor, we performed a permutation test on the mean of the average RNA expression from the ligand cluster and the mean of the ADT expression from the receptor cluster. Only ligand-receptor pairs with a *P*-value of lower than 0.05 were defined as significant pairs.

### Calculation of average and relative ranking of RNA and ADT expression

For the analysis of the ligand-receptor interactions identified through CiteFuse and the conventional approach using only mRNA expression, we calculated the concordance of mRNA and ADT expression of receptors. Because the same gene may be predicted to be involved as a receptor in a ligand-receptor interaction in multiple clusters, we performed a cluster-specific analysis as the expression and correlation of the mRNA and ADT of the receptor is likely to be different between clusters. Therefore, we evaluated concordance between mRNA and ADT in a cluster-specific and relative manner by calculating the ranking of mRNA and ADT expression in the cluster of interest in relation to all other clusters. We then plotted the relative ranking of mRNA and ADT expression against one another. For ligands, we also calculated a cluster-specific ranking based on their mRNA expression.

## Data and code availability

All data used in this study are available under accession numbers GSE126310. Sources for code used in this study are available from http://SydneyBioX.github.io/CiteFuse/.

## Author contributions

H.J.K. and Y.L. conceived the study with input from J.Y.H.Y. and P.Y.; H.J.K. and Y.L. developed the computational methods, tools, and the R package and led the data analysis and interpretation with input from J.Y.H.Y. and P.Y.; T.A.G. contributed to the development of computational methods; Y.L. implemented the Shiny app with input from H.J.K., J.H.Y.L. and P.Y.; H.J.K., Y.L., and P.Y. wrote the manuscript with input from J.Y.H.Y.; All authors revised, edited, and approved the final version of the manuscript.

## Acknowledgments

The authors thank their colleagues at the School of Mathematics and Statistics, The University of Sydney, for informative discussion and valuable feedback. This work is supported by an Australian Research Council (ARC)/Discovery Early Career Researcher Award (DE170100759) and a National Health and Medical Research Council (NHMRC) Investigator Grant (1173469) to P.Y., a National Health and Medical Research Council (NHMRC)/Career Development Fellowship (1105271) to J.Y.H.Y., an ARC/Discovery Project (DP170100654) grant to P.Y. and J.Y.H.Y., Research Training Program (RTP) and Chen Family Research Scholarship to Y.L, and Australian Research Council (ARC) Postgraduate Research Scholarship and Children’s Medical Research Institute Postgraduate Scholarship to H.J.K.

## Supplementary figures and legends

**Figure S1.**
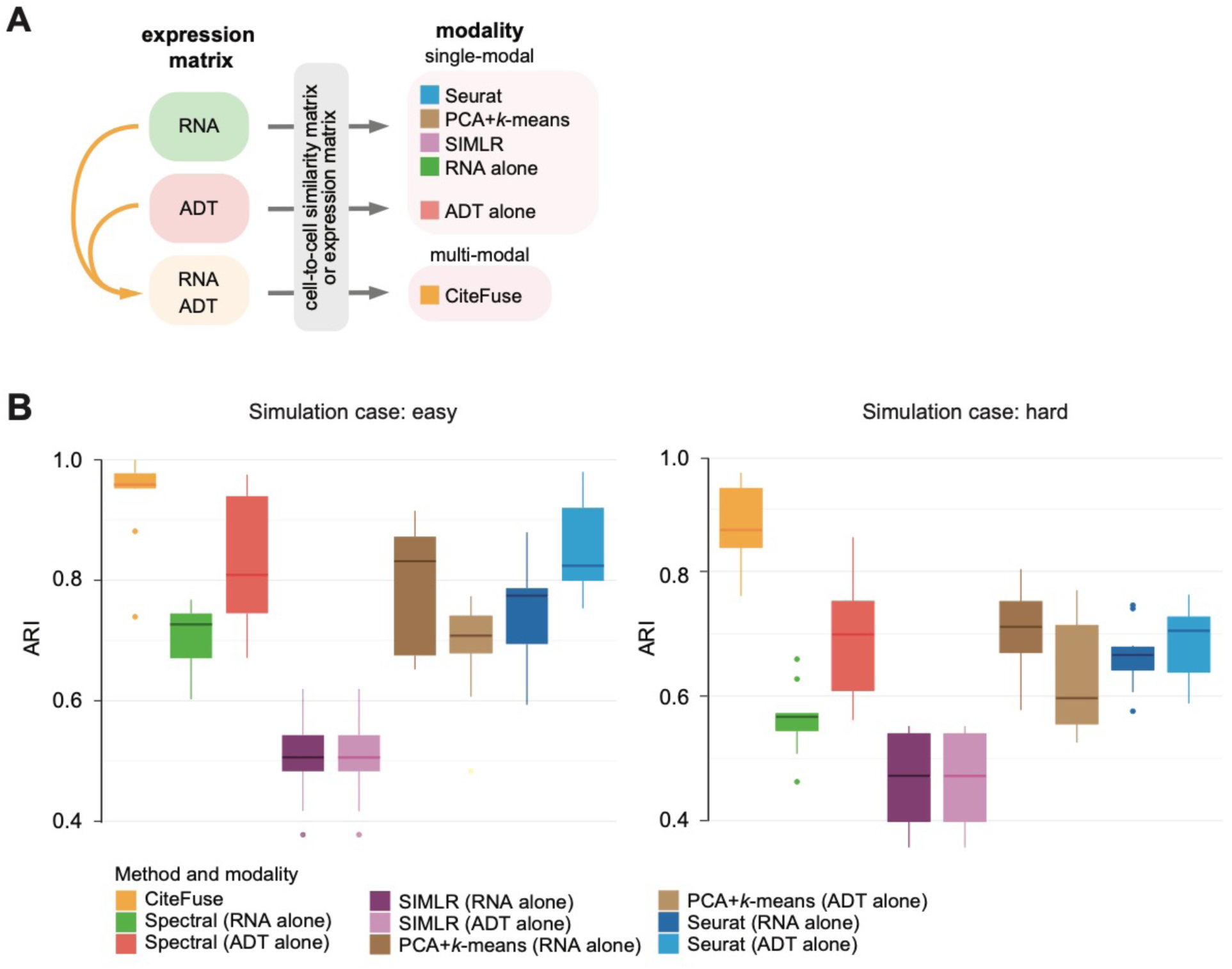
Evaluation of CiteFuse and other alternative methods using simulations (related to Figure 1). (A) A schematic summary of different methods and data modalities used for clustering cells. (B) Ten simulations were conducted for an easy and a hard scenario, respectively. Y-axis shows the adjusted rand index (ARI) calculated for clustering outputs from using various methods and data modalities on each of the two scenarios were presented as boxes.

**Figure S2.**
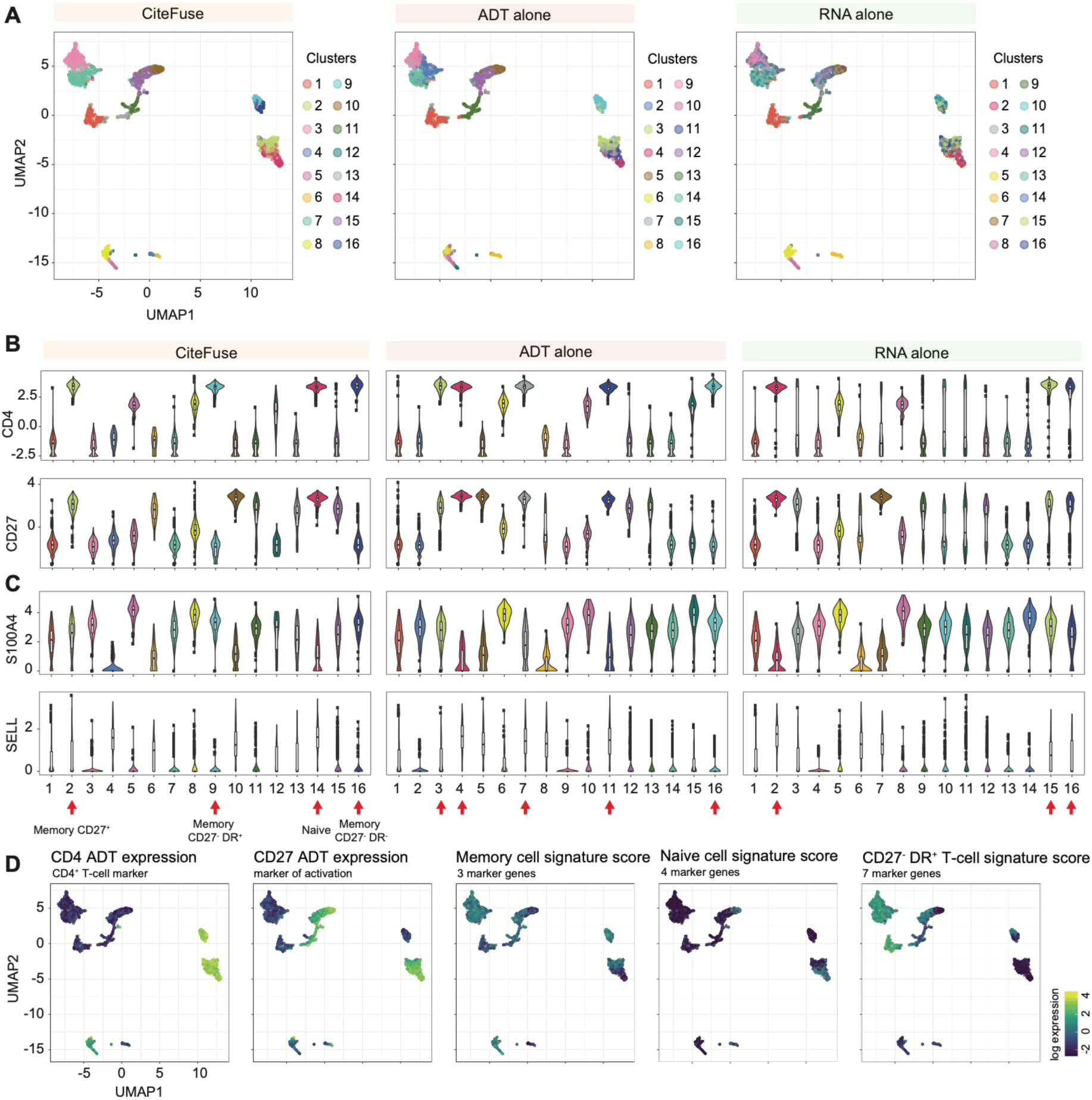
Clustering of CITE-seq data using single- or multi-modality (related to Figure 1). (A) UMAP of the fused expression matrix, ADT-alone and RNA-alone expression matrix of the human PBMC CITE-seq data (Mimitou et al., 2019). Clustering outcomes are highlighted by coloured points for both multi-modality (CiteFuse) and single-modality (ADT or RNA) approaches. (B) Centred log-ratio (CLR, y-axis) transformed ADT expression of CD4 and CD27 epitopes in clusters defined from CiteFuse, ADT-alone, and RNA-alone and (C) log RNA expression of S100A4, a marker of CD4+ memory T-cells, and SELL, a marker of naive CD4+ T-cells, in clusters defined from each approach. Clusters correspond to memory CD27+, CD27-DR+, CD27-DR-, and naive cells are highlighted by red arrows. (D) CLR-transformed expression of ADT (CD4 and CD27; first two panels) and log RNA expression of a set of signature genes for memory, naive, or CD27-HLA-DR+ CD4+ memory cells (third, fourth, and fifth panels) highlighted on UMAP of fused similarity matrix. A brighter colour denotes higher expression.

**Figure S3.**
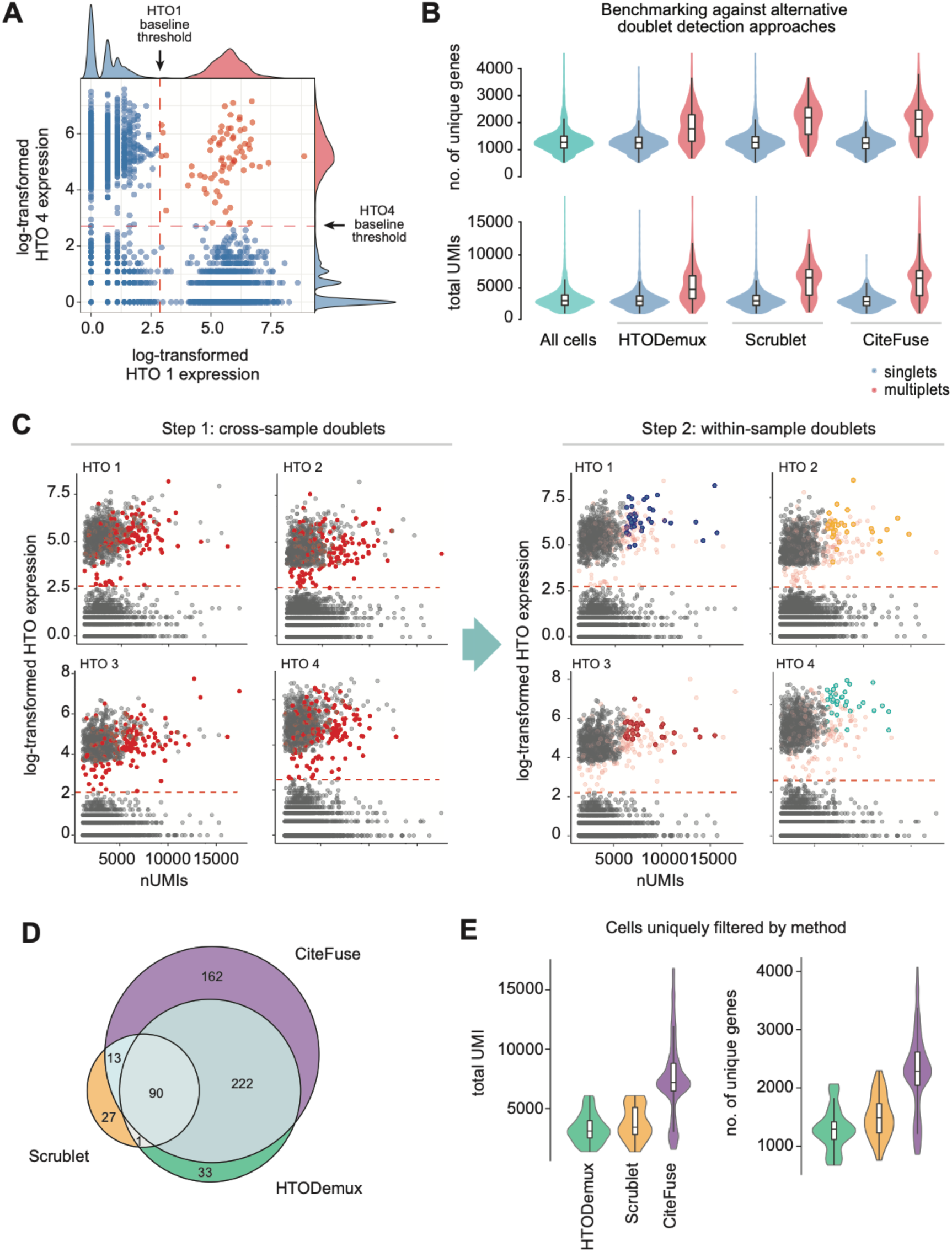
Cross-sample and within-sample doublet detection of CiteFuse (related to Figure 2). (A) Gaussian mixture modelling of log-transformed hashtag oligonucleotide (HTO) expression to identify cross-sample doublets (red points). (B) Total number of unique molecular identifiers (nUMI) and total number of genes expressed in all cells (both filtered and unfiltered) and HTODemux-,Scrublet-, and CiteFuse-identified singlets and doublets/multiplets. (C) A scatter plot of nUMI and log-transformed HTO expression for each HTO (1-4) highlighted by cross-sample doublets (red; left panel) and within-sample doublets (color-coded by HTO sample; right panel). (D) A Venn diagram of doublets depicting the overlap in identified doublets between the three filtering methods. (E) nUMI and total number of genes expressed in doublets uniquely identified by each filtering method.

**Figure S4.**
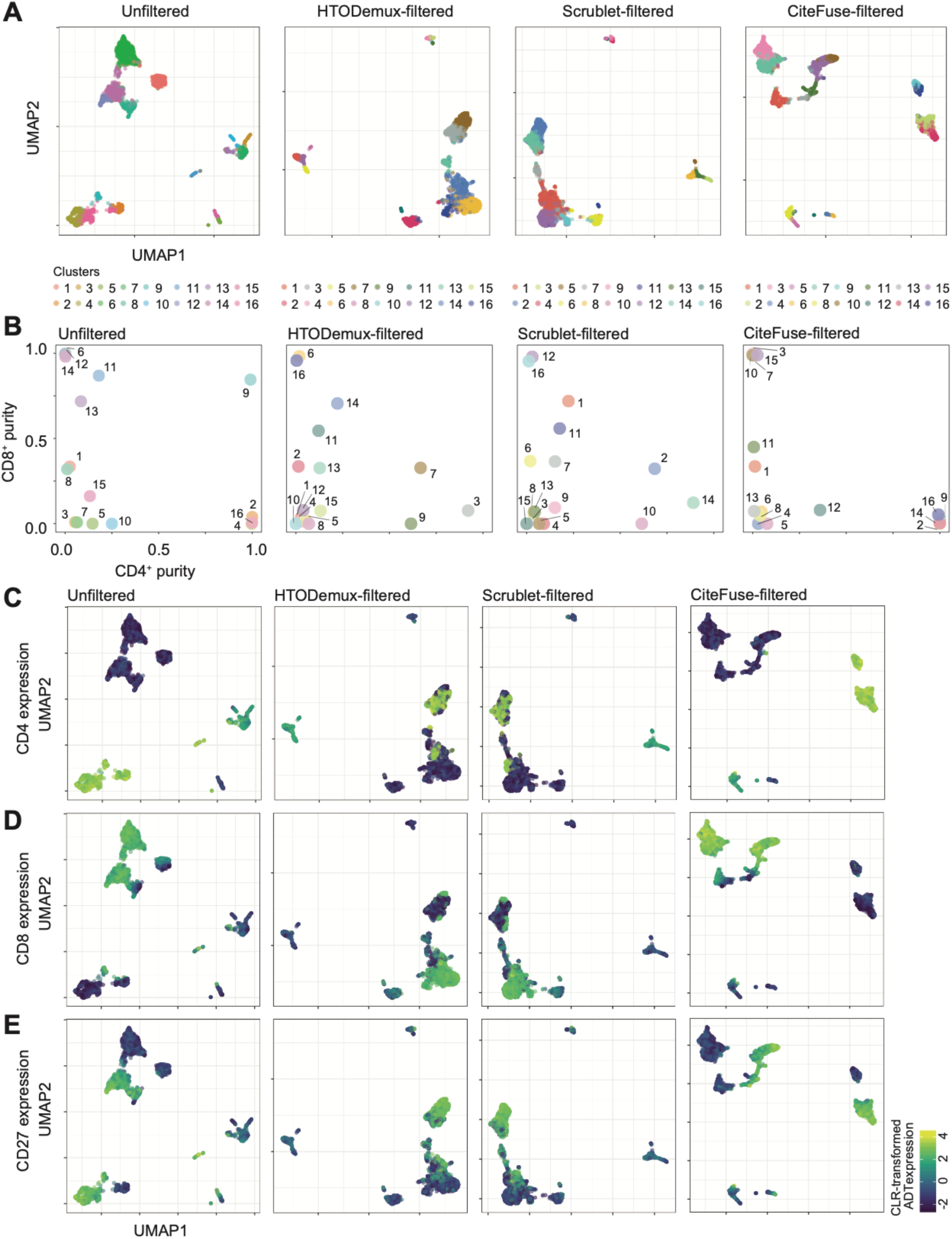
Clustering results from unfiltered and doublet filtered data (related to Figure 2). (A) UMAPs of the unfiltered, HTODemux-filtered, Scrublet-filtered, and CiteFuse-filtered matrix. Clusters generated by fused matrix of both unfiltered and filtered data are highlighted in different colours. (B) Purity scores of CD8+ cells (y-axis) against CD4+ (and CD11c-) (x-axis) cells in individual clusters by unfiltered data or data filtered by each of the three methods. CLR-transformed ADT expression of (C) CD4 (D) CD8 and (E) CD27 highlighted on UMAPs from (A).

**Figure S5.**
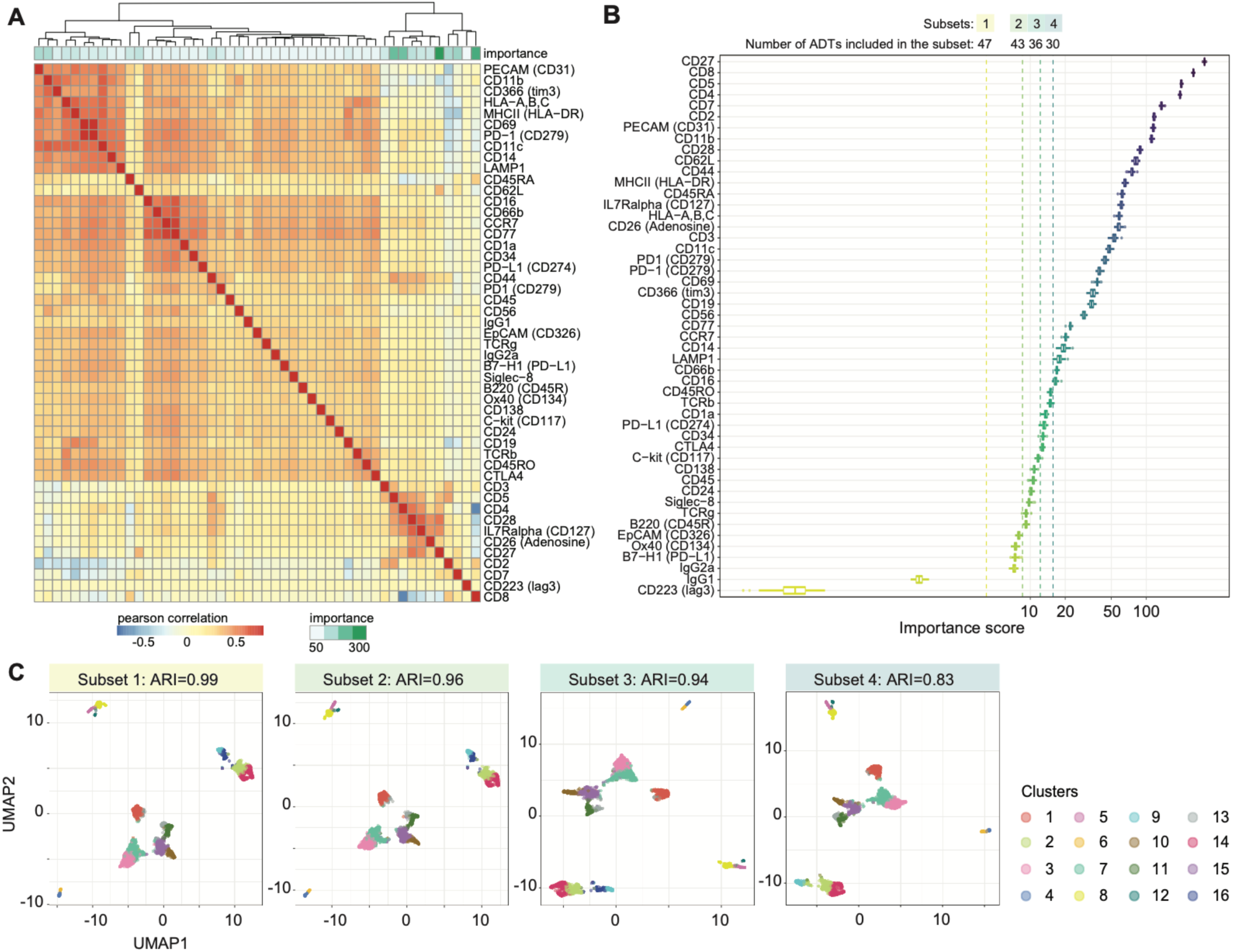
Evaluation of ADTs on CiteFuse clustering outputs (related to Figure 2). (A) Heatmap of pairwise correlation of ADT expression. Importance score of each ADT was generated by fitting a random forest on CiteFuse clustering outputs of fused matrix (see Methods). (B) Importance scores (x-axis) of ADT towards CiteFuse clustering outputs calculated as the average Gini index after 10 repeated fitting of random forest model. (C) UMAP of CiteFuse with various subsets of ADTs (in decreasing order from left to right panels) and adjusted rand index (ARI) of clustering outcomes against the full ADT set.

**Figure S6.**
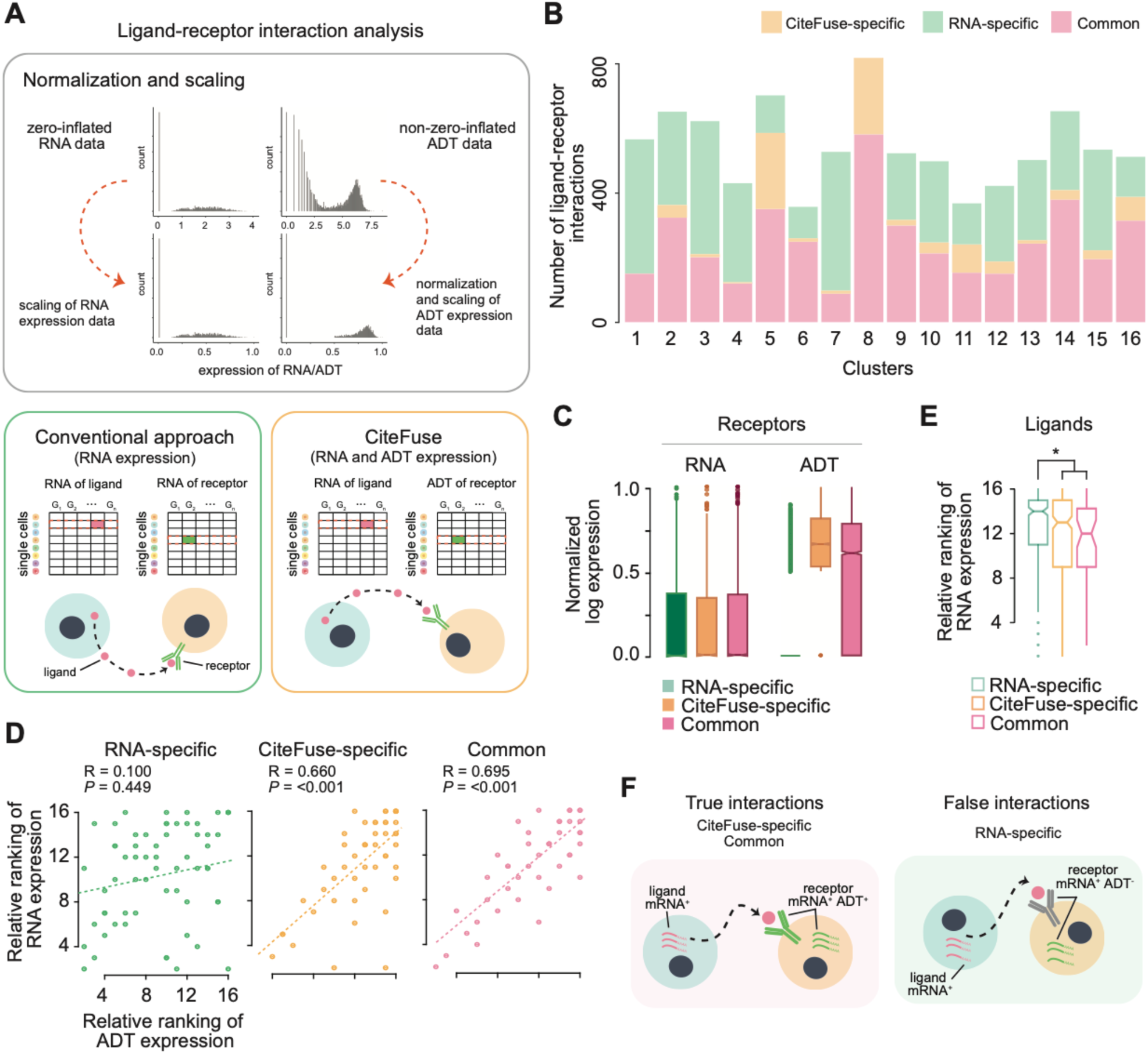
Ligand-receptor interaction prediction with CiteFuse (related to Figure 2). (A) A schematic illustrating the pre-processing step in CiteFuse (scaling of the RNA and ADT expression data and normalization of ADT expression data) and the two types of ligand-receptor interaction prediction methods: 1) conventional approach based on only RNA expression data and 2) CiteFuse approach based on both RNA and ADT expression to predict ligand-receptor interactions. (B) Number of ligand-receptor interactions predicted for each cluster by both conventional approach and CiteFuse (Common), or only by conventional approach (RNA-specific) or CiteFuse (CiteFuse-specific). (D) Scatter plot of relative ranking of RNA and ADT expression across clusters for all receptors identified in a ligand-receptor interaction in each of the three categories (i.e. RNA-specific, CiteFuse-specific, and Common). (E) Relative ranking of RNA expression of ligands in the three categories predicted by conventional approach and/or CiteFuse. (F) Schematic illustration of true and false ligand-receptor interactions and their mRNA and ADT expression where “+” and “-” denote high and low expression, respectively.

